# Default Mode Network in Temporal Lobe epilepsy: interactions with memory performance

**DOI:** 10.1101/205476

**Authors:** Tamires Araujo Zanão, Tátila Martins Lopes, Brunno Machado de Campos, Mateus Henrique Nogueira, Clarissa Lin Yasuda, Fernando Cendes

## Abstract

**Objective:** to investigate abnormal functional connectivity in the resting-state default mode network (DMN) and its relation to memory impairments in patients with temporal lobe epilepsy with and without hippocampal sclerosis (HS)

**Method:** we enrolled 122 MTLE patients divided into right-HS (n=42), left-HS (n=49), MRI-negative MTLE (n=31) and controls (n=69). All underwent resting-state seed-based connectivity fMRI, with a seed placed at the posterior cingulate cortex, an essential node for the DMN. In addition, patients and 41 controls were tested for verbal and visual memory, estimated intelligence coefficient and delayed recall.

**Results:** Both right-HS and MRI-negative group presented the poorest visual memory scores, and right-HS and left-HS had a worse performance in verbal memory compared to controls and MRI-negative groups. As expected, hippocampus was less connected than controls in all groups of patients. Although EEGs indicated that 64.5% of MRI-negative patients were lateralized to the left, this group showed activations similar to the right-HS.

**Conclusion:** Our data suggest that there is a disruption of the normal pattern of DMN in MTLE. Patients with left and right-HS presented similar, increased and decreased connectivity in the ipsilateral hemisphere; however, left-HS had abnormal decreased connectivity in the contralateral hemisphere. Per neuropsychological examination, the presence of HS in the left hemisphere had more impact on verbal memory, which was not found when the seizure focus is in the left hemisphere in the absence of HS. The absence of hippocampal atrophy seems to yield a less prominent disruption in both functional connectivity and neuropsychological performance.

## INTRODUCTION

Mesial temporal lobe epilepsy (MTLE) has been traditionally associated with memory impairment, especially with episodic memory, since hippocampus has a major involvement in consolidation of long-term memories ^1, 2^. However, it is also notable that MTLE patients frequently present more extensive cognitive damages not easily explained only by the temporal seizures focus ^3^. It is not surprising that recent studies have been investigating different brain networks in MTLE given the widespread, extra temporal structural and functional alterations observed in these patients, ^4, 5^. One particular network, the default mode network (DMN), has been focus of interest possibly because of a putative controversial recruitment of temporal lobe structures, including the hippocampus ^6^. DMN encompasses medial prefrontal and medial, lateral, and inferior parietal cortices, cerebellum and precuneus and, for some authors, the mesial temporal lobe is also included ^7^.

Regardless of controversies about hippocampal involvement ^7, 8^, several evidences indicate the impairment of DMN in MTLE ^9, 10^. The temporal lobe is vastly connected with the DMN; and hippocampus itself has both structural and functional connectivity to the posterior DMN ^7, 11^. Recent fMRI studies have revealed different patterns of spontaneous activity in specific nodes of the DMN in MTLE patients. While the general functional connectivity showed decreased activity within the DMN, areas as posterior cingulate cortex and right middle frontal cortex were hyper-connected compared with healthy subjects ^6, 12^. In addition, connectivity analyses with different techniques detected abnormal interactions between DMN and other networks in MTLE patients ^13–15^. Overall, given the important connections between hippocampus and key structures of DMN ^16^ it is tempting to speculate that the DMN impairment observed in MTLE could be associated with cognitive scathe of higher order brain functions.

Although some studies have disclosed DMN abnormalities in MTLE, most combined different underlying etiologies, failing to evaluate patients with and without hippocampal sclerosis (HS) separately. It is expected that patients with MTLE present episodic memory impairments (related to autobiographical memories) secondary to damaged structures directly involved in this cognitive function, such as hippocampus; however, it is still essential to better understand the extension of these damages and the influence it has in other cognitive aspects. In addition, quite few have analyzed the impact of the side of hippocampal atrophy on DMN alterations. Specifically, in MTLE associated with HS ^17^, side of hippocampal atrophy plays a key role in cognitive performance. It has been consistently accepted that side matters being the left hippocampus more related to verbal memory, while right hippocampus associates with nonverbal memory ^18, 19^. Hence, it is expected that left-HS patients present more verbal deficits while right-HS patients normally present more nonverbal deficits. However, scarce information exists about cognition and DMN in MRI-Negative MTLE patients and so far, there is no study that considered all these variables together.

In the present work, we used resting-state, seed-based connectivity method to analyze the DMN, combined with cognitive tests for memory and other domains. To identify the DMN, a seed was placed at the posterior cingulate cortex (PCC), which was previously described as an essential node for the DMN ^16, 20^. To investigate the impact of both side and presence of MRI-defined HS in the DMN and cognition, MTLE patients were separated into those with right-HS, left-HS, and MRI-negative (MTLE without HS). Our main hypothesis was that DMN would be severely disrupted in patients with hippocampal atrophy, followed by a more coherent network in the MRI-negative group; and that the pattern of this disruption would correlate with the neuropsychological assessment.

## MATERIALS AND METHODS

### Subjects

We included 122 MTLE patients (age range 21-70, 78 females, mean age of 46 years) divided into right-HS (n=42) and left-HS (n=49) and MRI-negative (n=31). They were recruited at the outpatient epilepsy clinic, University of Campinas - UNICAMP (Campinas - São Paulo, Brazil).

All patients had ictal semiology described by a close relative or documented by a medical staff during regular visits or during video-EEG monitoring. In addition, all had clinical and EEG features consistent with the diagnosis of MTLE according to the International League Against Epilepsy classification ^21^, and had no other findings suggesting an extra temporal focal epilepsy^22^. Also, 69 healthy controls (age range 23-66, 44 females, mean age of 44 years) were enrolled.

The HS was defined by visual MRI analysis and confirmed by hippocampal volumetry as described previously ^23^. HS signs were defined as hippocampal atrophy, hyperintense T2/FLAIR signal, and other MRI signs of HS, including loss of internal architecture in the hippocampus ^17^.

All subjects included are literate Brazilian Portuguese native speakers and were submitted to structural and functional brain imaging. In addition, all patients and a subset of 41 controls underwent neuropsychological assessment applied by neuropsychologists. The study was approved by the local ethical committee of UNICAMP and all participants signed informed consent forms.

## Interventions

### MRI data acquisitions

All images were acquired with a 3T Philips Intera Achieva (Philips, Best, the Netherlands) at the Neuroimaging Laboratory - LNI, University of Campinas - UNICAMP, with an eight-channel head coil.

#### 1) *Functional images*

Subjects were instructed to keep their eyes closed and to avoid goal-oriented thoughts. They underwent to an echo planar image (EPI) acquisition with isotropic voxels of 3 mm, axial plane with 40 slices, no gap, matrix ◻=◻80×80, flip angle ◻=◻90°, TR ◻=◻2s, TE ◻=◻30ms (6 min scan), resulting in 180 dynamics.

#### 2) *Structural images*

The MRI protocol included 3D-T1 weighted images (isotropic voxels of 1 mm for reconstruction in any plane, acquired in the sagittal plane; 180 slices, 1 mm thick, flip angle = 8°, TR=7.0 ms, TE=3.2 ms, matrix = 240 × 240), T2 weighted multi-eco coronal images (voxel sizes: 0.9×1×3 mm, TR = 3300 ms, TE=30/60/90/120/150 ms, matrix=200 × 180), as well as 3 mm thick coronal T1-inversion recovery perpendicular to the long axis of hippocampus (voxel sizes: 0.7×0.7×3 mm, TR = 3550 ms, TE=15 ms, IR delay = 400 ms, matrix=240 × 230) and FLAIR coronal and axial images (voxel sizes: 1×1×4 mm, TR = 11000 ms, TE=150 ms, IR delay = 2800 ms, matrix=232 × 192 for both plans).

### Image processing

Image preprocessing, analysis and statistical inferences were performed using the UF^2^C toolbox (http://www.lni.hc.unicamp.br/app/uf2c/) standard pipeline ^24^. The toolbox runs within MATLAB platform (2014b, The MathWorks, Inc. USA) with SPM12 (Statistical Parametric Mapping 12, http://www.fil.ion.ucl.ac.uk/spm/), as previously described ^24^.

### Seed based functional connectivity

The seed based method consists of selecting regions of interest (ROI) and correlating the time-series signals of the ROI with the rest of the brain. BOLD similarity and synchronicity between different gray matter areas indicate positive correlation ^25^, generating individual statistical maps. To obtain individual DMN maps (first-level analysis), a seed (1 cm^3^) was placed at the PCC (MNI coordinate: 0, −51, 15) (fig.1).

For the ROI time-series extraction, UF²C used the average time series of all ROI voxels that matched two consecutive criteria: i) being included in the subject GM mask and ii) if its correlation with the average time-series is higher than the average minus the standard deviation of all correlations between the ROI-masked voxels ^24^. This procedures warranty reliable functionally representative ROI average time-series.

To access group inferences (second-level analysis), SPM full-factorial model was used to compare patterns of activation for each group and between them. Six main comparisons were defined, between each group of patients and the control group (right-HS ×controls; controls × right-HS; left-HS × controls; controls ×left-HS; MRI-negative × controls and controls × MRI-negative). We also tried to correlate values from neuropsychological assessment with images but failed to detect significant results. All activations reported included clusters with at least 10 voxels and a threshold of p<0.001 uncorrected for multiple comparisons ^26^. The anatomical classification of SPM maps was accessed with *xjView toolbox (http://www.alivelearn.net/xjview)*, while images used for illustration were built with *MRIcroGL (www.mccauslandcenter.sc.edu/mricrogl/home)*.

### Neuropsychological evaluation

Neuropsychological assessment was selected to investigate different aspects of cognition, including verbal and visual memory, estimated intelligence coefficient (estIQ) and delayed recall (table e-1 – supplemental data).

#### Neuropsychological measures and analysis

EstIQ was measured using Vocabulary and Cubes sub-items of WAIS III ^27^, while others domains were evaluated through Wechsler Memory Scale Revised ^28^ and RAVLT ^29^. We used the Brazilian Portuguese validated version of all tests.

We performed a preliminary assumption testing to check normality, linearity, univariate and multivariate outliers, homogeneity of variance-covariance matrices and multicollinearity, with no serious violations noted. Given the identification of high collinearity between delayed recall and verbal memory (0.858) we performed a multivariate analysis (MANOVA) as first analysis with IQ, verbal memory, and visual memory; a separated univariate analysis with delayed recall was performed, as delayed recall reflects maintenance of information after time and interferences of goal oriented activities. Analyses were covaried for number of education years, as this variable differed between groups, and can interfere with cognitive fulfillment.

**Figure 1.**
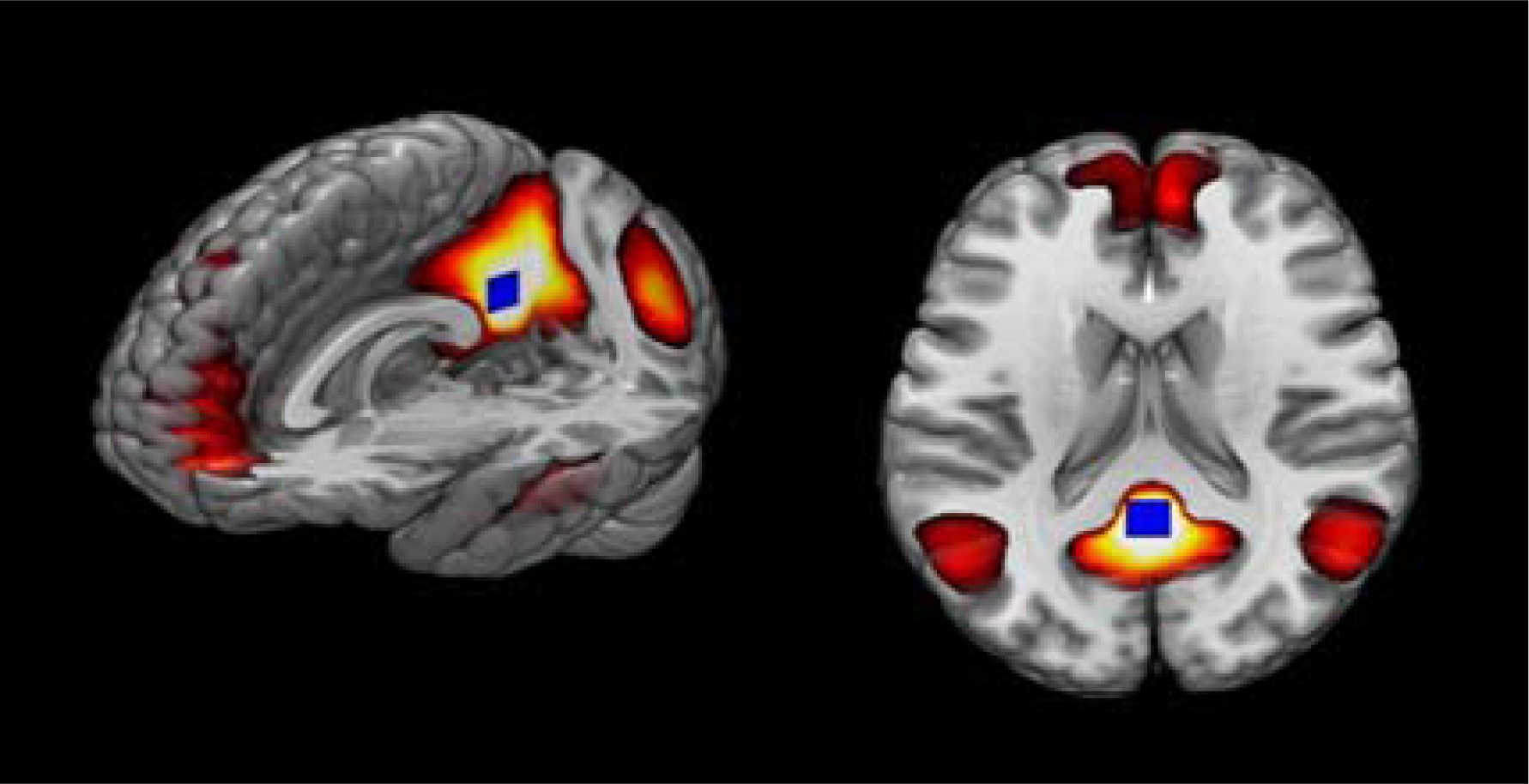
In blue, seed based placed at PCC and in yellow/red, the correlations of this ROI with others brain regions. The darker colors indicate stronger similarities in BOLD between PCC and the whole brain.

## RESULTS

### Demographic

Patients and controls were balanced for age and gender. However, left-HS group presented lower education, compared to controls (table 1). EEG investigation for MRI-Negative showed that 64.52% (20) of patients in this group were left sided, followed by 25.8% (8) undefined and only 9.68% (3) were right lateralized. EEG was lateralized to the side of hippocampal atrophy in all patients with HS.

### Neuropsychological assessment

Multivariate analyses (covaried for education years) revealed significant differences among groups, (F _(3, 428)_ = 3.57, p<0.01, Pillai`s Trace=0.56, partial η^2^ =0.56). Pairwise Bonferroni comparisons showed:

1. For verbal memory, MRI-negative group had similar performance to controls, while right-HS and left-HS showed poorer performance than controls (p<0.01) (Fig. 2-A).
2. For visual memory, right-HS and MRI-negative presented a worse performance than controls (both p<0.001); by contrast, left-HS did not differ from controls, although there was a tendency for lower scores in the left-HS (p=0.054). (Fig. 2-B)
3. For estIQ, all groups of patients presented worse accomplishment than controls (right-HS p=0.01, left-HS p<0.01, MRI-negative p=0.04). (Fig. 2-C)

Considering the importance of delayed recall (Fig 2-D) and the restriction to include this variable in the multivariate analysis, we performed a univariate analysis for delayed recall. Results showed worse performance for all patients’ groups compared to controls (right-HS p<0.01, left-HS p=0.01 and MRI-negative, p=0.02).

The images in Figures 3 (A, B and C) show the comparisons between each group of patients (right-HS, left-HS, and MRI-negative) with controls. Blue areas represent increased connectivity in patients, while red areas characterize reduced connectivity in patients, compared to controls.

Regardless side of HS, MTLE patients presented areas with similar patterns of increased connectivity with the DMN (seed on PCC), considering ipsilateral and contralateral hemispheres. In the ipsilateral hemispheres, frontal lobe was more connected, while temporal lobe, caudate and hippocampus presented reduced connectivity. The contralateral hemisphere also presented some similarities: both groups showed increased connectivity on parietal lobe associated with reduced connectivity on the temporal lobe. In addition to these common patterns, some peculiarities of connectivity with the DMN (seed on PCC) appeared relevant in contralateral hemisphere for each group in comparison to controls (table 2, table e-2):

1. Right-HS group also showed increased connectivity in frontal lobe, temporal lobe, and cerebellum; reduced connectivity in right-HS included limbic lobe
2. In left-HS, pallidum was more connected while reduced connectivity encompassed areas in both frontal lobe and insula.
3. In MRI-negative group, connectivity was increased bilaterally in frontal lobes and basal ganglia. Interestingly, reduced connectivity in the right hemisphere included the same areas (temporal lobe, caudate and hippocampus) identified in the ipsilateral hemisphere of right-HS and left-HS. While enhanced connectivity was identified in left hemisphere (frontal, temporal and parietal lobes), we did not identify reduced connectivity in the left hemisphere, compared to controls. Moreover, different from the other groups, increased connectivity on MRI-negative group was observed in basal ganglia bilaterally, right cerebellum and left occipital regions.

There were intriguing similarities between right-HS and MRI-negative (equivalent areas were less connected in the right hemisphere of both groups, and more connected regions were the same in left hemisphere for both). However, most patients in the MRI-negative group had left-sided EEG lateralization as described above. These results suggest that the presence of HS and seizure focus on the left side is associated with a different pattern of abnormal functional connectivity and memory dysfunction than left-sided EEG focus in MTLE without HS.

**Table 1.** Demographic and clinical information

|  | **Right HS** | **Left HS** | **MR-negative** | **Controls** | **P - value** |
| --- | --- | --- | --- | --- | --- |
| Mean SD or proportion |  |  |  |  |  |
| **Gender (female/male)** | 29/13 | 32/17 | 17/14 | 44/25 | 0.57 |
| **Age** | 47.1 ± 9.3 | 45.6 ± 9.8 | 46.9 ± 12.2 | 44.5 ± 11.3 | 0.12 |
| **School years** | 7.6 ± 4.4 | 7.3 ± 4.2 | 9.3 ± 4.4 | 10.2 ± 3.8 | <0.01 |
| **Duration of epilepsy (years)** | 30.3 ± 12.8 | 32.4 ± 12.2 | 27.8 ± 14.0 | n/a | 0.77 |
| **Total** | **42** | **49** | **31** | **69** |  |

Right-HS; right temporal lobe epilepsy, left-HS; left temporal lobe epilepsy, MR-Negative; magnetic resonance imaging negative, SD; standard deviation.

**Table 2.** Principal connected areas comparing patient groups with controls for ipsilateral and contralateral hemispheres

|  | **Ipsilateral** | **Contralateral** |
| --- | --- | --- |
| **Right-HS>controls** | Frontal Lobe | Frontal Lobe + Temporal Lobe + Parietal Lobe + Cerebellum |
| **Right-HS<controls** | Temporal Lobe + Caudate + Hippocampus | Temporal Lobe + Limbic Lobe |
| **Left-HS>controls** | Frontal Lobe | Parietal Lobe + Pallidum |
| **Left-HS<controls** | Temporal Lobe + Caudate + Hippocampus | Frontal Lobe + Temporal Lobe + Insula |
|  | **Right hemisphere** | **Left hemisphere** |
| **MR-Negative>controls** | Frontal Lobe + Basal Ganglia + Cerebellum | Frontal Lobe + Temporal Lobe + Parietal Lobe + Basal Ganglia + Occipital |
| **MR-Negative<controls** | Temporal lobe + Caudate + Hippocampus | --No activations-- |

Areas activated considering ipsilateral and contralateral sides for Right-HS and Left-HS and right and left hemisphere for MRI-negative group.

**Figure 2.**
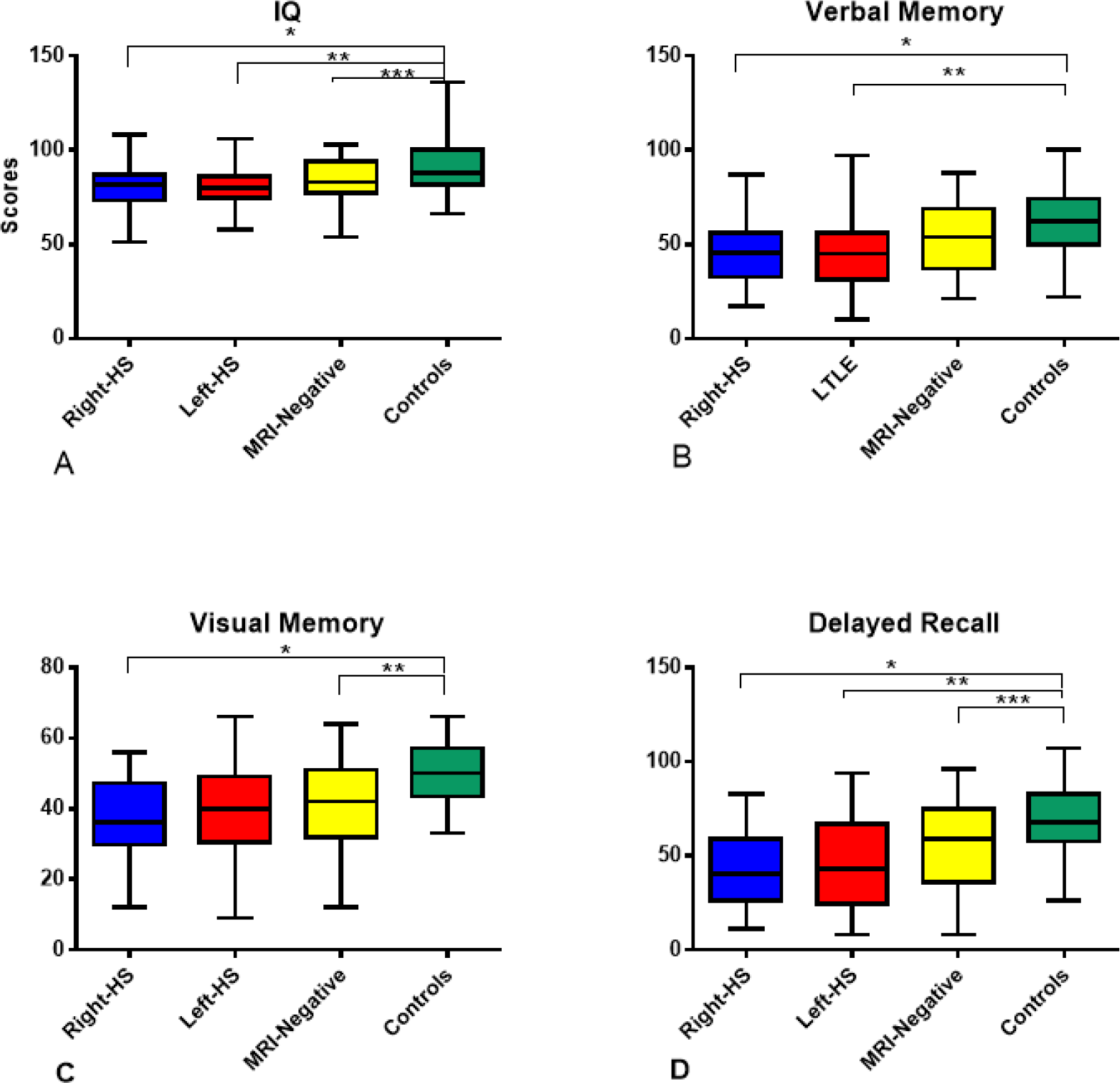
**A)** Comparison between right-HS, left-HS, MRI-negative and controls’ estIQ performance. * p=0.1, **p<0.01, ***p=0.04. B) Verbal memory comparison between groups. *p=0.01, **p=0.01. **C)** Comparison between right-HS, left-HS, MRI-negative and controls’ visual memory performance. *p=<0.01, **p<0.01. **C)**. **D)** Delayed recall comparison among groups. All patients’ groups presented a worse performance than controls *p<0.01, **p=0.01, ***p=0.02.

**Figure 3.**
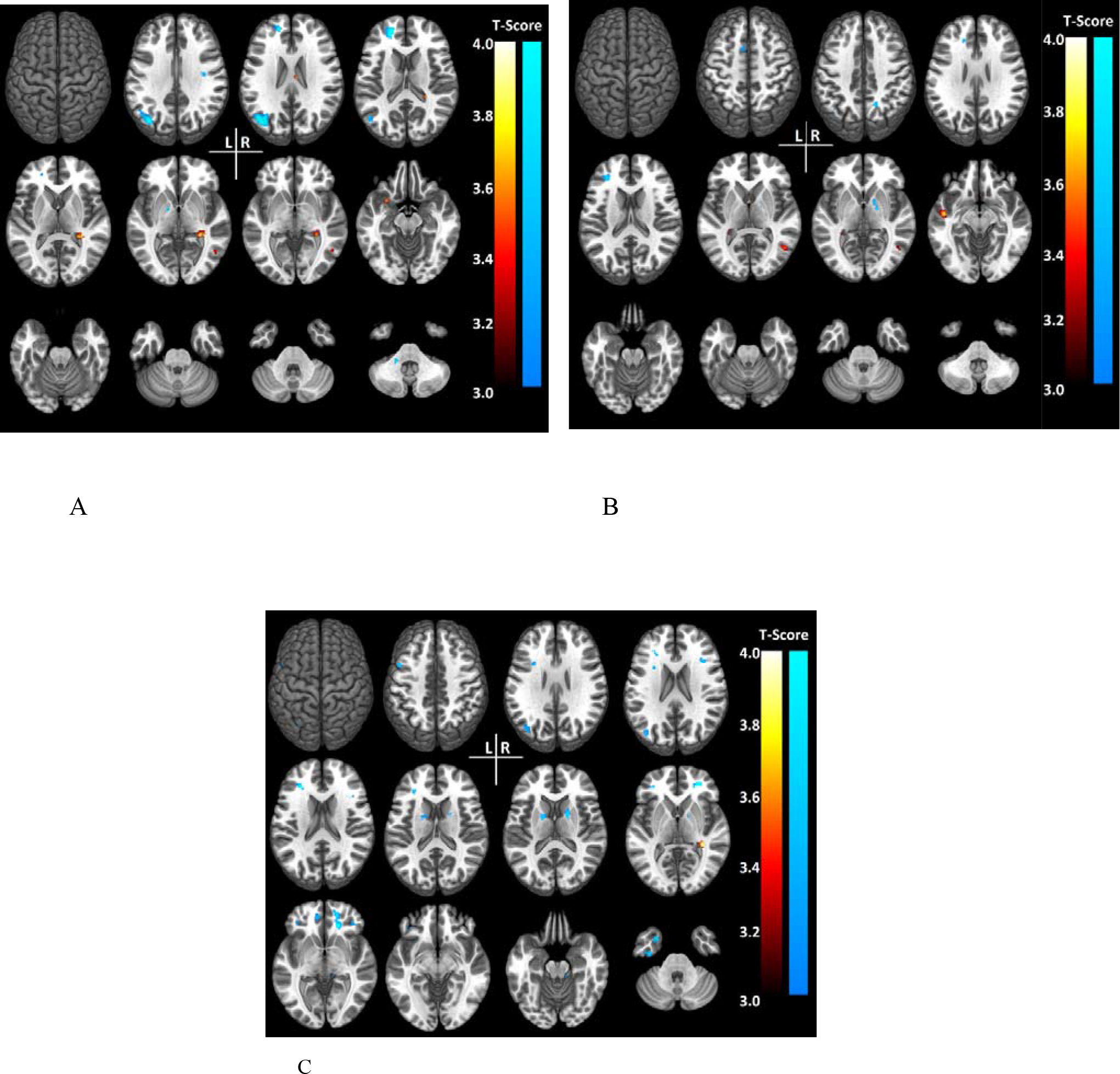
For all images, we represented in blue areas that are more connected in patients group than in controls and in red areas that are less connected in patients compared to controls (seed based on PCC). A - Right-HS compared to controls B - left-HS compared to controls and C - MRI-Neg compared to controls.

### DISCUSSION

Herein, we bring mutually key elements related to the DMN participation in MTLE cognitive impairments that, to our best knowledge, have not been entirely explored in previous studies. It is remarkable that regardless the presence or side of HS, MTLE patients presented an intricate pattern of increased and decreased connectivity, involving ipsilateral and contralateral hemispheres. Patients with left and right sided HS presented similar, more connected areas in ipsilateral hemisphere (mainly frontal lobe) associated with more connected areas in the contralateral parietal lobe. MRI-negative presented the same pattern as right-HS besides the left EEG lateralization in most of them.

Interestingly, reduced connections in temporal lobe, caudate and hippocampus were similarly identified in ipsilateral hemisphere of right-HS, left-HS and right hemisphere of MRI-negative. In addition to the commonalities, each group presented some particularities, such as increased connectivity in the basal ganglia of MRI-negative group, in the contralateral cerebellum of the right-HS and contralateral pallidum of left-HS. Furthermore, left-HS group had reduced connectivity in the contralateral frontal and temporal lobes, while the right-HS group had increased connectivity in these regions.

Not surprisingly, for MTLE, hippocampus is less connected than controls in all groups of patients. On the other hand, given the strong connections with hippocampus, the unanimous activation of frontal lobe in comparison to controls is newsworthy ^30^. Once neuroimaging and neuropsychological studies indicate the involvement of frontal and pre-frontal cortex in various aspects of memory, including learning, recall, remote retrieval, monitoring and controlling information processes ^31^, the recruitment of frontal lobe combined with memory impairment showed by neuropsychological assessment indicate that malfunction of connections, probably between frontal lobe and hippocampus, may be associated with disruptions on DMN. The over activation of frontal lobe seems to be a compensatory system to deactivation of hippocampus ^6^.

However, it is neither efficient nor enough to normalize cognitive performance. Regardless of frontal lobe recruitment, patients persist with frontal lobe dysfunction, suggesting that overall cognitive impairment may be associated with abnormal connections between hippocampus and frontal lobe ^32^. Studies indicate that cognitive impairment in MTLE are embracing and may occur outside the temporal lobe ^11^. Other authors have already considered that the main characterization of DMN dysfunction in MTLE patients consists in disrupted connections between the anterior and posterior regions ^12^. However, here we could characterize areas with more or less connections considering the presence and size of atrophy.

Relating to patients with HS, the results shown above may indicate that the presence or localization of atrophy is little or non-determinant for the ipsilateral hemisphere’s DMN connectivity. Considering recent studies that have revealed the importance of DMN for cognitive skills and memory ^33, 34^, the differences of performance between right-HS and left-HS may reside in the contralateral hemisphere (dys)connections and not in the damage of the ipsilateral mesial temporal lobe itself. Our results indicate some similarities between MRI-negative and right-HS groups in terms of connectivity, but the MRI-negative group presented more connections in both hemispheres when compared to other groups, besides the left EEG focus for most of our MRI-negative patients. Per neuropsychological examination, on contrary, both right-HS and left-HS had a worse performance in verbal memory compared to controls and MRI-negative. Little is known about MRI-negative patients and this result indicates that the presence of atrophy may be more harmful to verbal memory. Also, despite the impaired hippocampal activation, the recruitment of other areas may have been sufficient to yield a normal verbal memory performance in the MRI-negative group. Different from other groups, MRI-negative presented increased connectivity of basal ganglia, which may be explained by their involvement in DMN as an important role in cognitive flexibility. Also, the striatum (largest basal ganglia's structure) participation is related to interactions between different resting state networks, such as dorsal attention and frontoparietal networks ^35, 36^, suggesting the influence of these areas in higher cognition, especially in attention related activities. As verbal memory is a cognitive function that also depends on attention, the increased connectivity of basal ganglia may somehow explain the better performance of MRI-negative group in verbal memory, compared to other TLE groups. Future studies are necessary to clarify the role of basal ganglia in DMN, and its impact on MR-negative patients’ cognition.

Visual memory is usually attributed to right hemisphere and our results confirm a worse performance for right-HS compared to controls and left-HS, as expected ^37, 38^. Although visual memory is classically associated with right hemisphere, in the last decades studies have hypothesized that this subtype of memory has a more widespread and bilateral brain’s representation ^39^. MRI-negative and right-HS groups performed similarly in visual memory - therefore worse than left-HS and controls - which may be explained by the similar patterns of functional connectivity between MRI-negative and right-HS, as well as the extensive representation that may characterize the visual memory. As we have already mentioned, little is known about MRI-negative patients, particularly considering DMN and cognition and our new results can add important information about the topic. For both delayed recall and IQ, MTLE groups performed poorer than controls, but similarly among them. It may be explained by the more complex and diffuse nature of these cognitive functions and the widespread pattern of abnormal functional connectivity.

Some limitations of our work consisted in not considering the type and quantity of antiepileptic drugs in use during the neuropsychological evaluation and the MRI acquisition. Further studies may consider those variables and might contribute to a better understanding of the DMN and its relation to cognition in patients with MTLE.

### CONCLUSION

Our data suggest that MTLE disrupts the normal pattern of DMN, as we observed reduction of temporal lobe recruitment in patients, especially in the left-HS group. The absence of HS (MRI-negative) seems to yield a less prominent disruption in both functional connectivity and neuropsychological performance in verbal memory. For visual memory, both right-HS and MRI-negative groups presented the poorest scores and, interestingly, they have similar abnormal functional connectivity patterns.

## Acknowledgement

São Paulo Research Foundation (FAPESP)

